# Production of antibodies in egg whites of chickens

**DOI:** 10.1101/2021.04.18.440364

**Authors:** Angel Justiz Vaillant, Belkis Ferrer-Cosme

## Abstract

**Background:** IgM, which participates in the primary immune response, is the primary antibody in egg whites. There is scant information about the production of antibodies in egg whites. This study describes the preparation of antibodies against bacterial antigens.

**Methods:** Enzyme-linked immunosorbent assay (ELISA) was used to detect the presence of anti-egg white antibodies. The antibodies were purified using affinity chromatography. Statistical analyses were performed using SPSS version 22. Statistical significance was set at P<0.05.

**Results:** Large amounts of anti-protein A antibodies were produced in chicken egg whites. The generation of anti-SpA antibodies was demonstrated by affinity chromatography from 9 d post-immunization egg white samples. Inhibition of agglutination was observed in samples containing anti-SpA antibodies, and agglutination at the bottom of the wells was observed in the negative samples.

**Conclusion:** Anti-protein A antibodies (IgM) were produced in the egg whites of the immunized hens. Bacterial growth in blood agar plates was observed only in specimens plated with egg whites from pre-immunized birds. Protein A-affinity chromatography was helpful for the characterization of anti-protein A antibodies. Inhibition of these antibodies was observed *in vitro*.

## Introduction

Eggs are laid by animals of different species, including birds. Avian eggs consist of an egg shell, egg yolk, and egg white, which constitute the embryo from which a chick develops. From day zero, the avian egg contains a powerful immune system comprising antibodies against each foreign agent to which her mother (laying hen) has been in contact with. Egg yolk possesses a high titer of immunoglobulin Y (IgY), which is equivalent to immunoglobulin G (IgG) in mammalian species, and egg white contains immunoglobulin M (IgM) and immunoglobulin A (IgA) [1].

Purifying immunoglobulin M from the egg white of avian species is of interest as a source of immunoglobulins for oral immunization to prevent infections. The concentration of immunoglobulin M in egg whites was 0.15 2. Oral administration of hyper-immune eggs has been used to prevent bacterial infections in several animal species, such as dental caries caused by *Streptococci* species in rats [3] and *E. coli* infections in piglets [4]. Egg immunoglobulins have been used in the immunodiagnosis and therapy of several infectious diseases [5].

The egg white of birds has several proteins and polypeptides, which could be advantageous to human health. Yao et al. (1998) proposed that avidin may be a beneficial vehicle for transporting toxins, drugs, radioisotopes, or therapeutic genes to intraperitoneal tumors [6]. We aimed to provide information about the production of egg white immunoglobulins against staphylococcal protein-A and investigate the inhibition pattern of *Staphylococcus aureus* growth in pre-and post-immunized laying hens and its implications [7,8].

Protein A is the first bacterial immunoglobulin receptor to be described. It is displayed on the cell wall of Staphylococcus aureus and has been shown to react in a non-specific manner with human IgG in immunodiffusion tests. Protein A has a molecular weight of 42 kDa. It binds to the Fc fragment of IgG produced by several animal species, including dogs, rabbits, hamsters, monkeys, and others [9]. The native protein A consists of five domains. Of these, four show high structural homology, contain approximately 58 amino acids, and have the capacity to bind to the Fc region of IgG [9]. Structural changes following protein A binding to the Fc region of IgG have been studied by nuclear magnetic resonance and spectroscopy, which showed that the interaction involves the Z domain of protein A and does not involve helix unwinding. The protein A binding site is located at the interface CH2-CH3 of human and guinea pig IgGs [9].

In addition to the Fc gamma domains of IgG, protein A can interact with Fab domains. It mediates conventional antigen binding by immunoglobulin (Ig) heavy chains belonging to the VH3+ family [9]. Hillson et al. (1993) reported the structural basis for the interaction between protein A and VH3+ Ig molecules. The results demonstrated that among human IgM molecules, specificity for protein A was encoded by at least 11 different VH3 germline genes. The binding of protein A to human IgM is similar to that of bacterial superantigen binding to T cells [10].

The use of avians in antibody production results in a reduction in the use of laboratory animals used for this purpose. In addition, immunized chickens produce larger quantities of antibodies than rodents in the laboratory. The hens are farmyard animals and are, therefore, less expensive than laboratory animals, such as rabbits. Antibodies developed in birds recognize more epitopes in mammalian proteins. It is more advantageous to use chicken immunoglobulins in immunoassays, which detect mammalian proteins. This is especially true when the antigen is a highly conserved protein such as a hormone. Immunizing chicken with a protein induces the production of anti-anti-idiotypic antibodies that can recognize the original antigen [11].

## Materials and Methods

All chemical and biological reagents were commercially available (Sigma-Aldrich Co.).

### Production of antibodies against protein A in egg white from birds

Six-month-old healthy layer chickens (brown Leghorn) were injected at multiple sites on the breast with one mg of protein A in 0.5 ml complete Freund’s adjuvant on day zero, and one mg of the same protein in 0.5 ml incomplete Freund’s adjuvant on day 9. Eggs were collected from laying birds before and ten days after the immunization. Egg whites were manually separated [12] and stored at 20°C.

### ELISA for detection of antibodies against protein A

The 96-well polystyrene microplates were coated with 500 ng of protein A (Sigma-Aldrich) in coating buffer for four h at 37°C. The microplates were washed four times with buffer (PBS-Tween-20) and blocked with 3% non-fat dry milk in PBS (25 μl/well) for one h at room temperature. The microplates were washed four times. Samples were then added to 50 μl of a 1:50 dilution of egg white. After an incubation period of one h at RT, the microplates were washed four times, and 50 μl of the peroxidase-labeled protein-A conjugate at a dilution of 1:3000 was added. The microplates were then incubated for one h at RT and washed four times. Tetramethylbenzidine solution (50 μl) was added to the plates. After further incubation for 16 min, the reaction was stopped and analyzed using a microplate reader at a wavelength of 450 nm [13,14].

### Purification of anti-protein A antibodies from eggs whites of 9 days post-immunized birds by affinity chromatography

Egg white immunoglobulins (300 μl) were purified using a commercially available protein A affinity chromatography kit (PURE-1A, Sigma Aldrich Co.). The manufacturer’s instructions were followed, and the antibody concentration was set at 0.4. The isolated anti-protein A antibodies were stored at − -20°C [12,11].

### Inhibition of *S. aureus* growth by anti-protein A antibodies in egg whites of birds

The effects of purified anti-protein A antibodies on *S. aureus* isolates were investigated. The neutralizing ability of purified anti-protein A antibodies was studied as follows: One ml of brain heart infusion (BHI) broth was placed in 11 sterile test tubes. An equal volume of ten μl of purified anti-protein A was added at a concentration of 2.5 μg/μl. An inoculum of the ATCC *Staphylococcus aureus* strain (ATCC #33592) was made to 0.5 commercially prepared McFarland scale standards (1=300 × 106/ml bacteria concentration). Then, ten μl of the inoculum was poured serially from tubes 1 to 10. Tubes 11 and 12 were used as controls [7].

Tube preparations were plated on blood agar and incubated overnight at 35°C. The optical densities of the bacterial growth were assessed and plotted against the bacterial concentrations at various serial dilutions (1:10, 1:100, or 1:1000).

### Agglutination inhibition of protein A-bearing *Staphylococcus aureus* cells by purified anti-SpA antibodies *in vitro*

Serial dilutions of 25 μl at a concentration of 60 μg/μl of purified anti-protein A were added in duplicate to 96 wells microtiter plates containing 20 μl of protein A-bearing *S. aureus* cells and incubated for 1 h at RT. Inhibition of agglutination was observed in positive (+) samples (presenting anti-SpA antibodies), and agglutination was observed at the bottom of the negative samples. Commercially available human serum (Sigma-Aldrich) was used as the positive control.

### Statistical analysis

Statistically, SPSS version 22 was used. A P<0.05 was considered significant.

### Ethical approval

This study was approved by the ethical research committee of the University of West Indies. Mona Campus. Jamaica.

## Results and Discussion

Numerous photometric determinations were conducted to ascertain the cut-off point of the test that ended in a mean absorbance value of three-fold that of the negative controls. The cut-off point was set at 0.55. The sandwich ELISA showed a specificity and sensitivity of 98% and 96%, respectively. This ELISA was normalized by determining the optimal reaction conditions and optimal coating reagent, antigen, conjugate, and substrate concentrations using checkerboard titration.

The use of protein A-coated microplates and peroxidase-labeled protein A conjugate ensured that the anti-SpA immunoglobulins could be tested positive in the sandwich ELISA because other bird antibodies do not bind to staphylococcal protein A in a non-specific manner. The ELISA showed a considerable concentration of anti-protein A antibodies (5 higher than the concentration in eggs of the pre-immunized birds) in the egg whites from birds immunized with the bacterial protein. One hundred% of the birds tested positive for anti-protein A antibodies, as shown in Table 1.

**Table 1:** Sandwich Enzyme-Linked Immunosorbent Assay (ELISA) for detection of anti-protein A antibodies in birds.

| Variables | Mean OD at 450 nm | SD |
| --- | --- | --- |
| Laying hen 1 | 1.308 | 0.053 |
| Laying hen 2 | 1.455 | 0.045 |
| Laying hen 3 | 1.387 | 0.050 |
| Laying hen 4 | 1.309 | 0.069 |
| Laying hen 5 | 1.415 | 0.071 |
| Laying hen 6 | 1.125 | 0.058 |
| Positive control 1 | 1.650 | 0.047 |
| Positive control 2 | 1.575 | 0.066 |
| Negative control 1 | 0.237 | 0.006 |
| Negative control 2 | 0.195 | 0.009 |
| Blank 1 | 0.088 | 0.003 |
| Blank 2 | 0.073 | 0.002 |

Anti-protein A antibody production was determined by affinity chromatography from nine days post-immunization egg whites [16]. When the bacterial concentration was serially diluted at 1:10, as shown in Table 2, the mean optical density of the bacterial growth in pooled egg whites of post-immunized birds was 1.475 versus pre-immunized birds that showed a mean OD value of 0.186. There was a notable difference in the absorbance values of pre-and post-immunized animals in the three serial bacterial dilutions. Comparable results were reported in an earlier study on the hindrance of the growth of *B. burgdorferi* in an *in vitro* testing of dogs for specific antibodies to the surface antigen [7].

**Table 2:** Showing the laboratory results of bacteria growth observed inimmunized and pre-immunized birds 9 days post-immunization. ^A^Bacteria concentration in ml after serial dilutions; ^B^optical density values of pooled egg white from post-immunized and pre-immunized birds; P values <0.05 are statistically significant.

| Bacterial concentration<sup>A</sup> | Immunized/pre-immunized hens<sup>B</sup> | P-value |
| --- | --- | --- |
| 1:10 | 0.186/1.475 | < 0.05 |
| 1:100 | 0.175/1.490 | < 0.05 |
| 1:1000 | 0.155/1.338 | < 0.05 |

The mixtures were streaked on a blood agar plate and incubated overnight, indicating growth inhibition of *S. aureus* in the plates incubated with anti-SpA antibody samples. Bacterial growth occurred only in the negative anti-protein A antibody specimens from pre-immunized birds. As shown in Table 3, the number of inhibited samples from post-immunized fowl was 47 out of 50, denoting a percentage inhibition of 96%, which was statistically significant. In pre-immunized fowl, the inhibition percentage was not statistically significant; the representation of inhibited samples was 4 out of 50, and it was considered that this number of samples, which inhibited the growth of *Staphylococcus aureus* in a non-specific mode, might be related to the presence of natural antibodies in the egg white specimens.

**Table 3:** Percentage inhibition of the growth of *S. aureus* in blood agarplates streaked out with egg white preparations. The presence of anti-SpA antibodies in the egg white was responsible for the inhibition of the growth of the bacterium *S. aureus*.

| Birds | N. of inhibited/Total | % Inhibition | P-value |
| --- | --- | --- | --- |
| Immunized laying hens | 47/50 | 94 | < 0.05 |
| Pre-immunized laying hens | 4/50 | 7 | > 0.05 |

Another outcome was the inhibition of agglutination of protein A-bearing *Staphylococcus aureus* cells by purified anti-protein A antibodies *in vitro*, which confirmed the prior findings. This could be explained by the coupling of anti-protein A immunoglobulins to SpA-bearing *S. aureus* cells, blocking those from agglutination at the base of the wells. A pooled positive specimen for anti-protein A antibodies in egg white samples hindered the agglutination of protein A-bearing *Staphylococcus aureus* cells at dilutions of 1:4096, in contrast to a pooled negative specimen that agglutinated protein A-bearing *Staphylococcus aureus* from 1:16 to 1:8192 dilutions.

The production of antibodies in egg whites of birds immunized with a protein has been reported by other authors, as Kowalczyk et al. (2019) reported that IgM was found only in egg white extracts. In comparison to IgY, IgM antibodies were not transferred to the serum of turkey poults [17]. Hamal et al. (2006) documented that chicks first synthesized IgM, followed by IgA and IgY. Anti-Newcastle disease virus (NDV) and anti-infectious bronchitis (IBV) antibody levels were detected in the plasma, egg yolks, and plasma of chicks on days 3 and 7 [18]. These results suggest the mechanism of transfer of IgM from the egg white to the chick serum and emphasize the importance of IgM in the bird’s immune system to protect against primary infections against bacteria, viruses, fungi, and chemicals.

## Conclusion

Anti-protein A antibodies (IgM) were produced in the egg whites of the immunized hens. Bacterial growth in blood agar plates was observed only in specimens plated with egg whites from pre-immunized birds. Protein A-affinity chromatography was helpful for the characterization of anti-protein A antibodies. Inhibition of these antibodies was observed *in vitro*.

